# Archaeal ribosomal proteins possess nuclear localization signal-type motifs: implications for the origin of the cell nucleus

**DOI:** 10.1101/671966

**Authors:** Sergey Melnikov, Hui-Si Kwok, Kasidet Manakongtreecheep, Antonia van den Elzen, Carson C. Thoreen, Dieter Söll

## Abstract

Eukaryotic cells are divided into the nucleus and the cytosol, and, to enter the nucleus, proteins typically possess short signal sequences, known as nuclear localization signals (NLSs). Although NLSs have long been considered as features unique to eukaryotic proteins, we show here that similar or identical protein segments are present in ribosomal proteins from the Archaea. Specifically, the ribosomal proteins uL3, uL15, uL18, and uS12 possess NLS-type motifs that are conserved across all major branches of the Archaea, including the most ancient groups *Microarchaeota* and *Diapherotrites*, pointing to the ancient origin of NLS-type motifs in the Archaea. Furthermore, by using fluorescence microscopy, we show that the archaeal NLS-type motifs can functionally substitute eukaryotic NLSs and direct the transport of ribosomal proteins into the nuclei of human cells. Collectively, these findings illustrate that the origin of NLSs preceded the origin of the cell nucleus, suggesting that the initial function of NLSs was not related to intracellular trafficking. Overall, our study reveals rare evolutionary intermediates among archaeal cells that can help elucidate the sequence of events that led to the origin of the eukaryotic cell.

## Introduction

Our understanding of past evolutionary events mainly relies on the discovery of transitional forms. Classical examples of these transitional forms include the fossilized bird-like dinosaur *Archaeopteryx lithographica* and the crawling fish *Tiktaalik roseae*, whose bone structures illuminated many aspects of how, between the Late Jurassic and the Late Devonian, the fins of ancestral fish were transformed into the legs of terrestrial animals and subsequently into the feathered wings of birds^1-3^. However, the further we delve into the past, the more we find ourselves limited in our archaeological record, so that the most ancient events in history of life on Earth are known in sketchy outline or remain enigmatic.

One such enigmatic event is related to the question how the cell nucleus originated. Fossil records indicate that, between 1.7 and 2.7 billion years ago, a group of ancestral prokaryotic cells were transformed into what we now refer to as eukaryotes: they acquired a DNA-storage compartment – the nucleus – that was separated from the cytoplasm by a nuclear membrane equipped with selectively penetrable channels, the nuclear pores^4,5^. To help them pass via the nuclear pores from the cytoplasm into the nucleus, cellular proteins have evolved specialized signal sequences, the nuclear localization signals (NLSs)^6^. A typical NLS is a short and surface-exposed stretch of basic residues that is recognized by specialized receptors, karyopherins^7^. Karyopherins transfer NLS-containing proteins across the nuclear pores^7^. Although the nuclear–cytoplasmic trafficking machinery includes more than 100 different components, including karyopherins, nuclear pore components, and other proteins, the order in which these proteins evolved and the factors that drove the transition from prokaryotic to eukaryotic cell structures are currently unknown. In the absence of evidence of a single transitional form between prokaryotes and eukaryotes, there are at least twelve alternative hypotheses of how the system of nuclear–cytoplasmic trafficking might have emerged^8^.

Previous studies have shown that many aspects of the early evolution of life on Earth can be understood through analysis of the most ancient macromolecules in a living cell, such as ribosomes. Because ribosomes are likely as old as life itself, their structure has been used as a living molecular fossil to gain an understanding of such evolutionary enigmas as the origin of catalytic RNA^9,10^, the evolution of protein folding^11,12^, the origin of the genetic code^13,14^, and the reason for the stereospecific structure of proteins^15^, among others^16-18^. Additionally, eukaryotic ribosomes have been shown to be adapted to the nuclear–cytoplasmic separation of eukaryotic cells because eukaryotic ribosomal proteins, unlike their bacterial counterparts, have evolved NLSs that allow ribosomal proteins to enter the cell nucleus where they are subsequently incorporated into nascent ribosomes^19^. This finding suggested that further investigation into ribosomes’ structures may lead to a better understanding of the origin of the nuclear–cytoplasmic trafficking^19^.

In this study we have analyzed ribosome structures from the three domains of life to investigate origins of nuclear localization signals in eukaryotic proteins. By comparing homologous ribosomal proteins from each of the three domains of life, we found that protein segments that were described as NLSs in eukaryotic ribosomal proteins were also present in homologous proteins from Archaea. This finding indicates that at least some NLSs evolved in proteins considerably earlier than the event when cells separated into the nucleus and the cytoplasm. This finding reveals a group of rare evolutionary intermediates – NLS-type motifs in archaeal ribosomal proteins, – which may shed light on the sequence of events that eventually resulted in the origin of the nuclear–cytoplasmic separation of eukaryotic cells.

## Results

### Archaeal ribosomal proteins possess nuclear localization signal-motifs

Seeking to better understand when NLS-motifs might have emerged in ribosomal proteins, we assessed their conservation among ribosomal proteins from the three domains of life. To date, NLS-motifs have been characterized in ten ribosomal proteins from several eukaryotic species (**Table 1**). We assessed the conservation of these NLS-motifs by using multiple sequence alignments of eukaryotic ribosomal proteins (from 482 species) and their homologs in the Bacteria (2,951 species) and Archaea (402 species) (**Supplementary Data 1**).

**Table 1.** Previously determined nuclear localization signals in eukaryotic ribosomal proteins.

| Protein | Old name | Organism | Sequence | Ref | Tag used to monitor the nucleolar accumulation | Presence in Archaea |
| --- | --- | --- | --- | --- | --- | --- |
| uS3 | S3 | *S. cerevisiae*<br>*H. sapiens* | 3-KKRK-7 | [49] | eGFP at the C-terminus<br>(*S. cerevisiae*) | - |
| uS4 | S9 | *H. sapiens* | 171-GRVKRKNAKKGQ-182 | [50] | eGFP at the N- and C-termini | - |
| uS8 | S22 | *S. cerevisiae* | 20-GKRQVLIRP-28 | [51] | β-galactosidase at the N-terminus | - |
| uS12 | S23 | *H. sapiens* | 2-<br>GKCRGLRTARKLRSHRRDQ<br>KWHDKQYKKAHLGTALKA<br>NPF-41 | [19] | eGFP at the C-terminus | + |
| uL3 | L3 | *S. cerevisiae* | 2-<br>SHRKYEAPRHGHLGFLPRKR<br>A-21 | [52] | β-galactosidase at the C-terminus | + |
| uL13 | L13a | *H. sapiens* | 59-RRK-61 | [53] | HA-tag, position is not specified | - |
| uL15 | L29 | *S. cerevisiae* | 6-KTRKHRG-13<br>23-KHRKHPG-29 | [54] | β-galactosidase at the C-terminus | + |
| uL18 | L5 | *H. sapiens*<br>*X. laevis* | 21-RRRREGKTDYYARKRLV-<br>37<br>255-KKPKKEVKKKR-265 | [55] | eGFP at the C-terminus | + |
| uL23 | L23a,<br>L25 | *S. cerevisiae*,<br>*H. sapiens* | 11-KKAVVKG-17<br>18-TNGKKALKVRT-28 | [56] | β-galactosidase at the C-terminus<br>(*S. cerevisiae*) | - |
| uL29 | L35 | *H. sapiens* | 71-<br>KGKKYQPKDLRAKKTRALR<br>RALTKF-95 | [57] | GST at the C-terminus | - |

We found that six NLS-motifs – including those in ribosomal proteins uS3, uS4, uS8, uL13, uL23, and uL29 – are highly conserved among the Eukarya but absent in Bacteria and Archaea (**Table 1**). This finding was consistent with studies of other proteins, such as histones, illustrating that NLSs are found only eukaryotic but not prokaryotic protein homologs^20,21^. However, four proteins – uL3, uL15, uL18, and uS12 – were found to have NLS-type motifs not only in the Eukarya but also in the Archaea (**Fig. 1, Supplementary Data 1**).

**Fig. 1.**
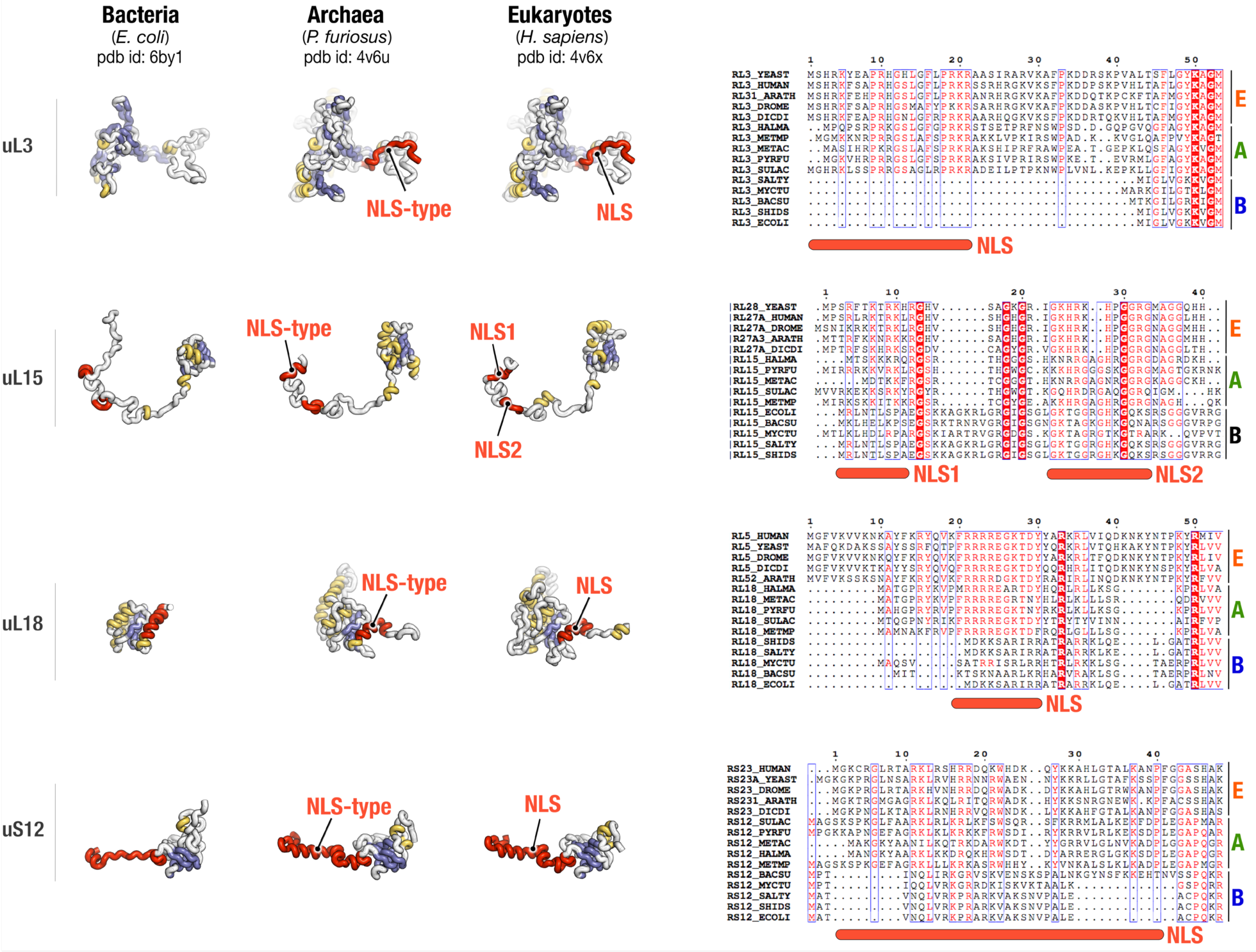
Archaeal ribosomal proteins have segments that are similar or identical to nuclear localization signals (NLSs). This figure compares the crystal structures and sequences of homologous ribosomal proteins from Archaea, Bacteria, and Eukarya. NLSs in eukaryotic proteins and corresponding segments in homologous archaeal proteins are highlighted in red. This figure illustrates that NLSs of eukaryotic proteins are absent in homologous bacterial ribosomal proteins, but present in homologous archaeal ribosomal proteins.

The ribosomal structure has been determined for several archaeal species, including *Haloarcula marismortui*^22^ and *Pyrococcus furiosus*^23^, so we therefore next investigated whether archaeal and eukaryotic NLS-type motifs have conserved structures. The NLS-type motifs in proteins uL3, uL15, uL18, and uS12 turned out to have conserved secondary and tertiary structures between Eukarya and Archaea (**Fig. 1**). Thus, our analysis revealed that NLS-motifs are not limited to eukaryotic proteins but can also be found in their archaeal homologs.

### NLS-type motifs are conserved in all groups of Archaea, including the most ancient archaeal branches

Having found NLSs-type segments in archaeal ribosomal proteins, we next asked whether these motifs are preserved in all branches of the Archaea or just in a subset of archaeal species. To answer this, we analyzed the conservation of NLS-motifs in the archaeal proteins uL3, uL15, uL18, and uS12 (**Fig. 2, Supplementary Data 2**). To date, archaeal species have been divided into four large lineages, including DPANN, Euryarchaeota, TACK and Asgard superphyla (**Fig. 2**)^24^. We anticipated that NLS-type motifs would be present only in the most recently evolved and eukaryote-like branches of archaeal species, such as Asgard. Indeed, we found that sequence similarity between eukaryotic NLSs and archaeal NLS-type motifs increases with the transition from ancient archaeal branches (DPANN) to more recently emerged branches (Asgard) (**Supplementary Data 2**). However, even in the DPANN superphylum the sequence similarity remains above 50% for each of the four NLS-type motifs (**Fig. 2**), with some DPANN species (e.g., *Mancarchaeum acidiphilum, Nanoarchaeota archaeon*, and *Aenigmarchaeota archaeon*) having only a single substitution in their NLS-type motifs (typically lysine-to-arginine) compared with eukaryotic ribosomal proteins (**Supplementary Data 2**). Thus, contrary to our expectations, we found that NLS-type motifs are conserved across all the archaeal branches, including the most ancient superphylum, DPANN.

**Fig. 2.**
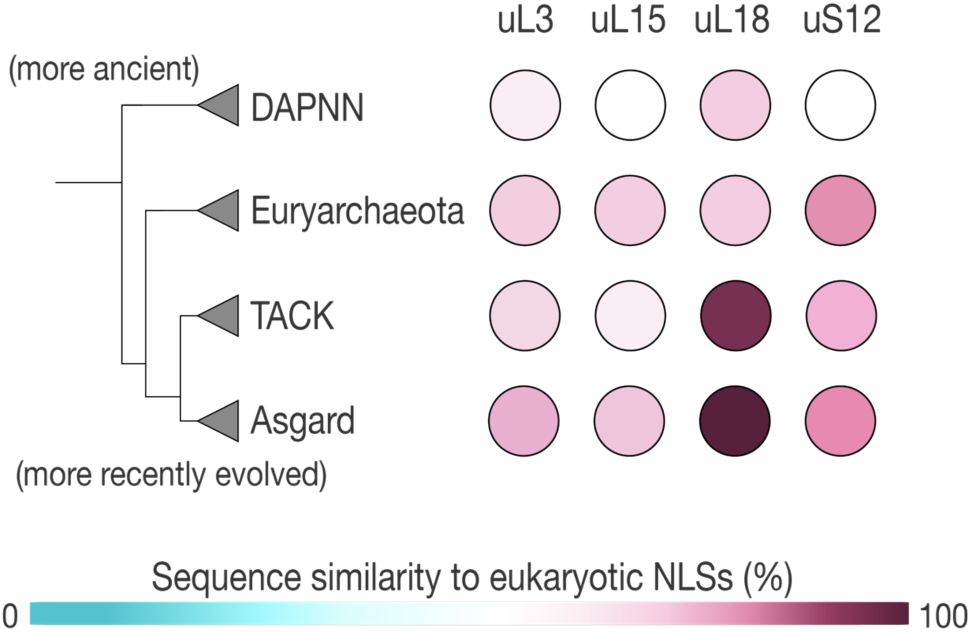
The NLS-type motifs are conserved in ribosomal proteins from all known branches of the Archaea. A phylogenetic tree of life, as according to (Ref. 24), showing the major branches of archaeal species and a diagram illustrating high sequence-conservation among NLS-type motifs in archaeal ribosomal proteins of the major groups of archaeal species, from the most ancient branches (DAPNN superphylum) to the most recently emerged eukaryote-like branch (Asgard superphylum).

Notably, many archaeal NLS-type motifs have precisely same sets of hydrophobic and basic residues as the NLS-motifs of eukaryotic ribosomal proteins, which is particularly common among species from the eukaryote-like Asgard superphylum (**Supplementary Data 2**). For instance, the NLS of the ribosomal protein uL18 in the eukaryote *Saccharomyces cerevisiae* has a set of hydrophobic and basic residues identical to those in 89 archaeal species, including species from *Pyrococcus* (Euryarchaeota), *Acidianus* (TACK), and *Lokiarchaeum* (Asgard) genera (**Supplementary Data 2**). Similarly, an identical set of hydrophobic and basic residues can be found in the NLS of *S. cerevisiae* protein uL3 and in the corresponding NLS-type sequences from 27 archaeal species (**Supplementary Data 2**). Furthermore, in 3 of these archaeal species uL3 has a preserved adjacent serine residue (Ser24 in *S. cerevisiae*), phosphorylation of which in yeast is thought to regulate uL3 intracellular transport and ribosome biogenesis^25^. Overall, this analysis revealed that even the most ancient lineages of archaeal species carry highly conserved protein segments that closely resemble eukaryotic NLSs.

### NLS-type motifs have evolved independently of changes in ribosomal RNA

Finding NLS-type motifs in archaeal proteins was surprising as it raised the question: why do organisms that lack a cell nucleus have conserved NLS-type motifs? What advantage would be conferred by having these signal sequences in archaeal cells? Many segments in ribosomal proteins have previously been shown to have coevolved with novel segments in rRNA^26-29^, so we sought to determine whether the emergence of NLSs in ribosomal proteins was somehow related to the evolution of ribosomal RNA. To answer this question, we analyzed NLSs’ contacts with rRNA (**Fig. 3**).

**Fig. 3.**
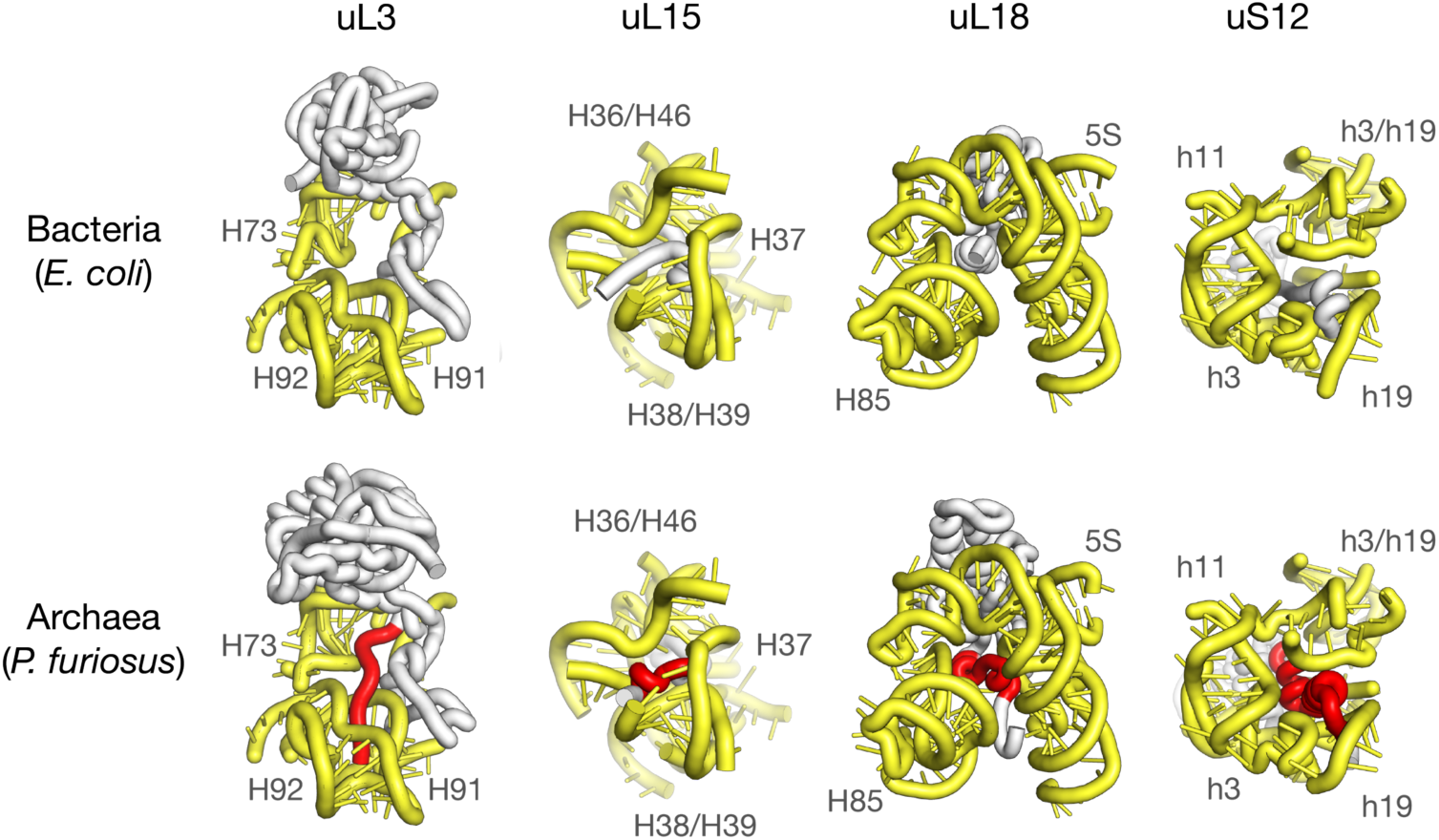
NLS-type motifs have emerged independently of changes in ribosomal RNA. The panels show the interior of bacterial (*E. coli*, PDB ID: 6by1) and archaeal (*P. furiosus*, PDB ID: 4v6u) ribosomes where the NLS-type motifs of ribosomal proteins (highlighted in red) and corresponding segments in bacterial ribosomal proteins interact with rRNA. The figure illustrates that Archaea/Eukarya-specific NLS-type motifs are buried in the ribosome interior where they bind highly conserved rRNA segments. In the archaeal ribosome structure, the NLS-type motifs bind rRNA helical junctions, suggesting that the NLS-motifs may help to recognize rRNA or govern rRNA-folding during ribosome biogenesis. Overall, this figure illustrates that the rRNA structure before and after the emergence of NLS-type motifs remains conserved, illustrating that NLS-type motifs in archaeal ribosomal proteins have evolved independently of changes in rRNA.

Previously, we showed that NLSs in ribosomal proteins are buried within the ribosome interior, and explained how these signals become inactivated during ribosome biogenesis to prevent nuclear import of mature ribosomes^19^. We also showed that, within the ribosomal structure, NLSs of ribosomal proteins interact with helical junctions, suggesting that NLSs may facilitate rRNA-folding during ribosome biogenesis^19^. Here, comparing ribosome structures from the archaeon *P. furiosus* and the bacterium *E. coli*, we found that the NLS-type motifs of archaeal ribosomal proteins bind highly conserved rRNA segments that have conserved tertiary structure in Archaea and Bacteria (**Fig. 3**). Thus, our analysis revealed that NLSs emerged in ribosomal proteins independently of changes in ribosomal RNA.

### Archaeal NLS-type motifs can substitute eukaryotic NLSs to direct intracellular transport of ribosomal proteins in eukaryotic cells

Finally, we explored whether archaeal NLS-type motifs can functionally substitute NLSs of eukaryotic ribosomal proteins. To test this, we replaced the NLS-type sequence in human ribosomal protein uS12 with the corresponding NLS-type segment of archaeal uS12 from *Sulfolobus sulfotaricus* or *Thermoplasma acidophilum* **(Fig. 4)**. As shown previously, deletion of the NLS impaires uS12 accumulation in the nucleoli, a site of ribosome biogenesis in the cell nucleus (although the truncated protein remains predominantly nuclear likely due the residual NLS activity of its globular domain)^19^. However, chimeric uS12 proteins carrying archaeal NLS-type motifs were localized in the nucleoli in a similar way to the wild-type human uS12 (**Fig. 4**). Thus, we found that NLS-type motifs of archaeal ribosomal proteins are not only similar in sequence and structure to eukaryotic NLSs, but can also complement the activity of eukaryotic NLSs.

**Fig. 4.**
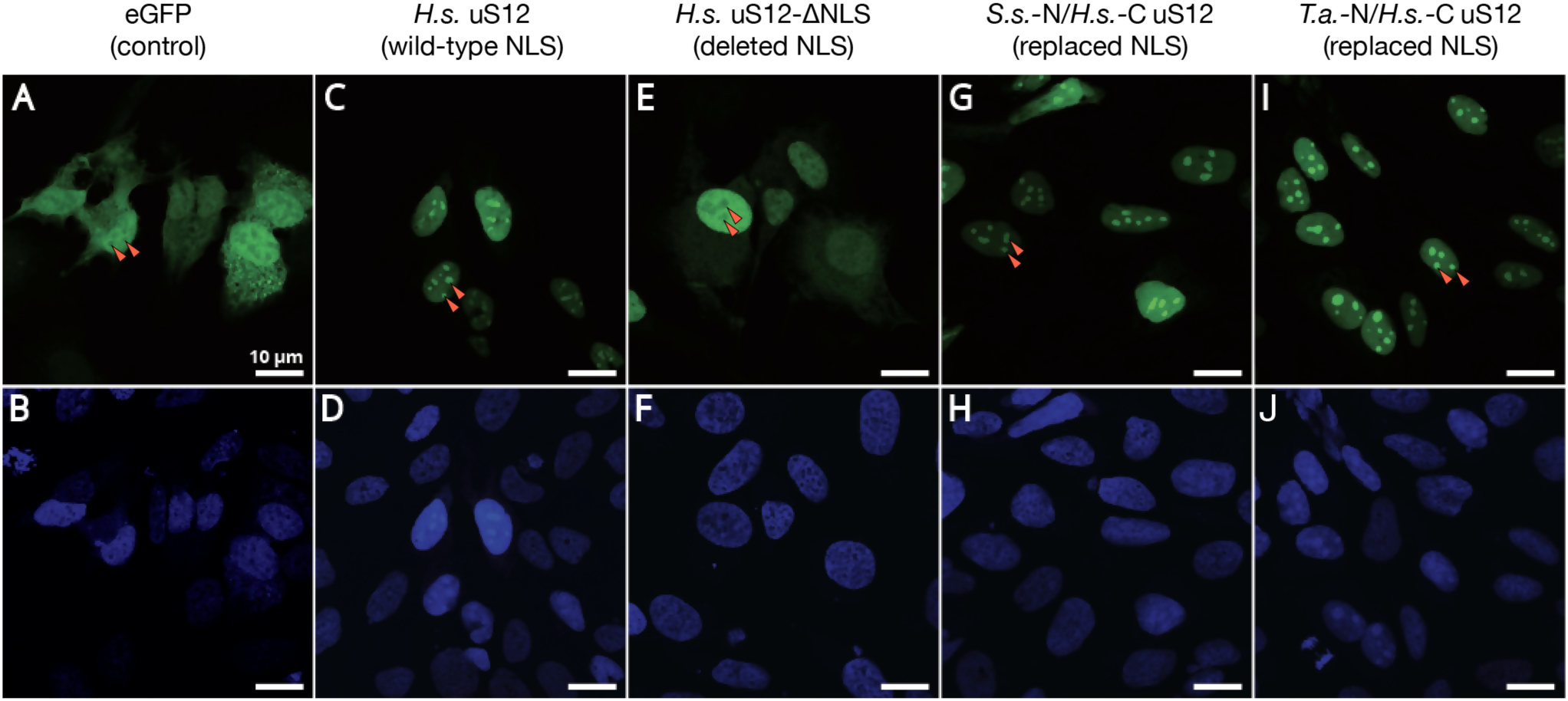
Archaeal NLS-type motifs can functionally substitute NLS-signals of the eukaryotic ribosomal protein uS12 in human cells. The panels show microscopic snapshots of eGFP fluorescence (green, top panels) and fluorescence of the DNA-staining agent DAPI (blue, bottom panels) in the human cell line HEK293T. Red arrows point to the location of human cell nucleoli. Cells are expressing the following eGFP fusions: (**A**,**B**) eGFP alone (as a negative control) shows both cytoplasmic and nuclear localization; (**C**,**D**) eGFP fusion with human uS12 accumulates in nucleoli; (**E**,**F**) eGFP fusion with human uS12 lacking the NLS-containing N-terminus (residues 1–41) is no longer localized in cell nucleoli; (**G**,**H**) eGFP fusion with human uS12 in which the N-terminus (residues 1–41) is replaced by the N-terminus of uS12 from the archaeon *Sulfolobus solfataricus*; (**I**,**J**) eGFP fusion with human uS12 in which the N-terminus (residues 1–41) is replaced by the N-terminus of uS12 from the archaeon *Thermoplasma acidophilum*. The panels **G**–**J** illustrate that N-terminal segments of the archaeal ribosomal protein uS12 (either from *S. solfataricus* or *T. acidophilum*) are able to functionally substitute the NLS-containing N-terminal segment of human uS12, to direct uS12 accumulation in human cell nucleoli.

## Discussion

In this study, we have identified a group of rare evolutionary intermediates that can shed light on the sequence of events that led to the emergence of the nuclear–cytoplasmic transport system. Our finding of NLS-type sequences in the ribosomal proteins uS12, uL3, uL15, and uL18 suggests that at least some cellular proteins evolved NLS-motifs long before the nucleus emerged, indicating that the initial function of NLSs could have been distinct from directing protein transport. Why the NLS-motifs emerged in ribosomal proteins in the Archaea – in species that have neither cell nucleus nor karyopherin proteins to recognize the NLS-type motifs – remains unclear^30,31^, as does what advantage might have been conferred by having these signals in the absence of nuclear–cytoplasmic transport. Furthermore, the initial function of NLS-type sequences in cells in the absence of the nuclear envelope and the nuclear–cytoplasmic trafficking system is unknown.

These questions resonate with several classical studies in evolutionary biology, such as studies on the origin of birds^32^. In the origin of birds theory, there is a famous question “of what use is half a wing?”. This refers to the fact that it took multiple millions of years to transform the limbs of ancestral reptiles into the wings of ancestral birds. Yet, for most of this time, the transitional forms of a wing could not support flight. Thus, the advantage of flight could not drive the positive selection of species with “germinal”, “premature” wings. Thus, what was the advantage of “half a wing”? Studies of modern birds suggest that wings could initially have emerged to facilitate gliding or accelerated climbing – two forms of motion related to flying that are preserved in the behavior of modern birds^33-35^.

Similar to the origin of wings, it is possible that NLSs could initially have emerged to fulfil similar, yet not identical, or incomplete functions of modern NLSs. In this regard, it is important to note that NLSs are known to fulfil three biological activities: (i) they participate in the recognition of nucleic acids, because most known NLSs reside within DNA- or RNA-binding protein domains^36^; (ii) they mediate protein transport across nuclear pores by recruiting karyopherins^6,7^; and (iii) karyopherin-binding to NLSs shields their electrostatic charges and prevents non-specific interactions of nucleic acid-binding domains^37^. In other words, karyopherins that bind NLSs not only fulfil the role of transport factors but also serve as chaperones for highly-charged proteins in eukaryotic cells^37^.

Based on our findings, it is tempting to suggest an order in which these three biological activities of NLSs may have evolved. High conservation of NLS-type motifs in archaeal ribosomal proteins and their abundant contacts with rRNA suggest that, at least in some proteins, the NLS-type sequences could have evolved as nucleic acid-binding domains that were required to increase the specificity of interactions in the increasingly complex macromolecular content of evolving cells (**Fig. 5a**). Later, these cells could have evolved a DNA/RNA-mimicking chaperone that would recognize NLS-type sequences in the way chaperones for hydrophobic proteins recognize hydrophobic protein segments to prevent non-specific interactions and protein aggregation (**Fig. 5b**). Only later, when this chaperone for charged proteins had evolved the ability to bind to the nuclear pores and direct protein transport to the nucleus, did the NLSs eventually become signals of intracellular transport (**Fig. 5c**). If this hypothetical order of events is correct, then the system of nuclear– cytoplasmic trafficking could originally have evolved as a system of chaperones for highly charged proteins, to prevent nucleic acid-binding domains from non-specific interactions in the increasingly complex environment of ancestral eukaryotic cells.

**Fig. 5.**
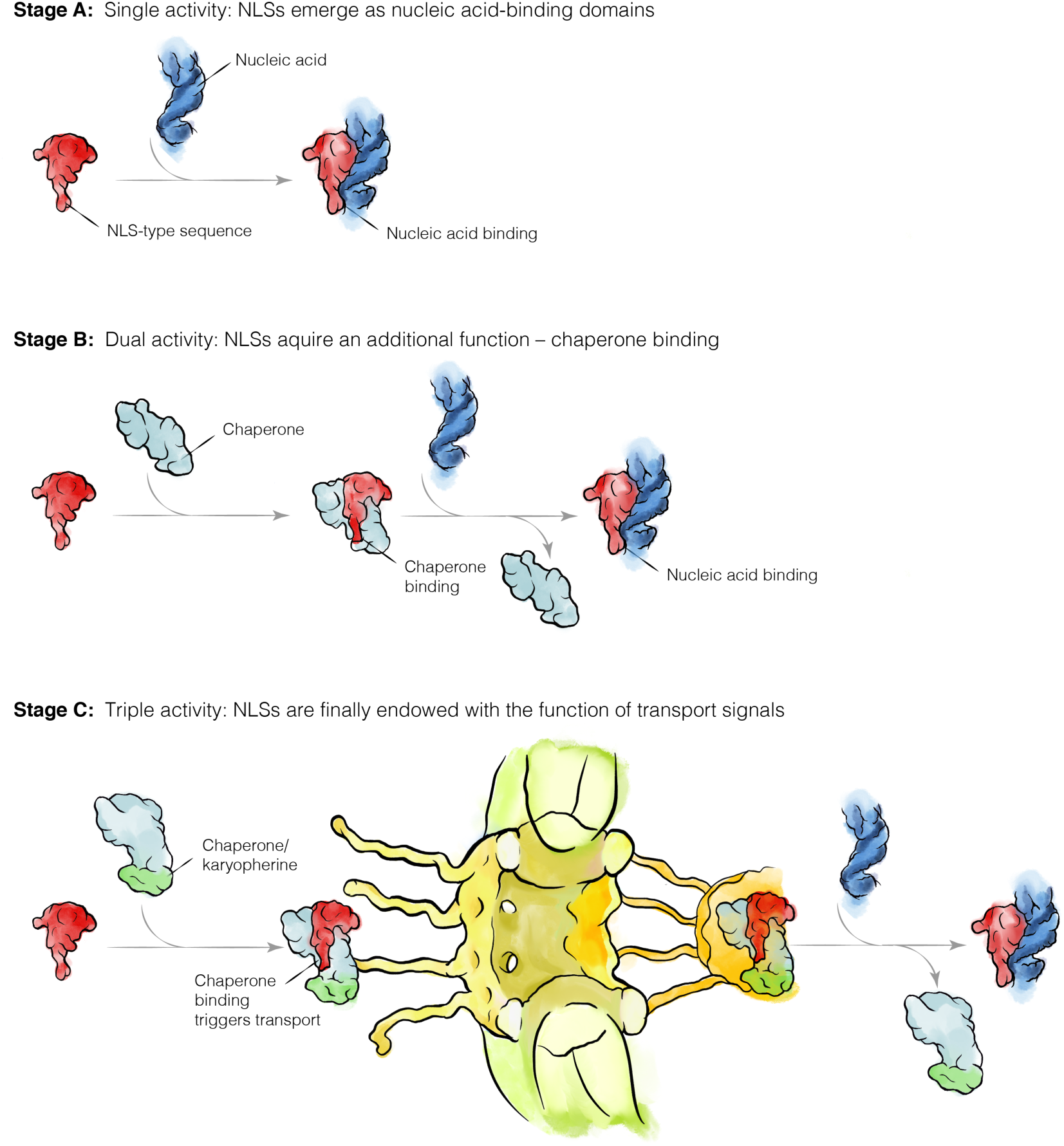
A hypothetical order of NLS evolution in eukaryotic proteins. In the modern eukaryotic cell, NLSs fulfil three biological activities: (i) they serve as signal peptides to direct protein transport into the nucleus; (ii) during this protein transport, they recruit trafficking factors (karyopherins) that shield high positive charge of nucleic acid-binding domains (which prevents non-specific interactions of NLS-containing proteins with other molecules in a cell); and (iii) typically, NLSs reside within DNA- or RNA-binding domains of proteins and, therefore, NLSs mediate specific recognition of nucleic acids. Our model suggests that NLSs may have originally emerged as nucleic acid-binding domains in a cell lacking the nuclear-cytoplasmic separation (**Stage A**). Later, a chaperone emerged that shielded the highly positively-charged nucleic acid-binding domains (**Stage B**). Finally, when cells got separated into the nucleus and the cytoplasm, this chaperone turned into a karyopherin when it acquired the additional capacity to mediate the long-distance trafficking of proteins across nuclear pores (**Stage C**).

Our study leaves unanswered the question: why do the non-globular extensions of ribosomal proteins have different structures in Bacteria and Archaea, even though these extensions bind invariable, universally conserved rRNA segments? Previously, it was suggested that these structural differences stemmed from an independent evolutionary origin for non-globular segments in bacterial and eukaryotic ribosomal proteins^11^. Another possible reason may be related to differences in ribosome biogenesis pathways between Bacteria (in which ribosome biogenesis largely occurs via self-assembly and with the help of protein-guided rRNA modification machinery) and Archaea/Eukarya (in which ribosome biogenesis requires archaea/eukaryote-specific biogenesis factors and RNA-guided machinery for rRNA modification)^38-41^. We also cannot exclude the possibility that these differences may be related not to the problem of ribosome evolution but to a more general problem of specific macromolecular interactions in a cell. It is not possible, based on our data, to exclude the notion that these non-globular protein segments were remodeled in the Archaea because the original non-globular segments in bacterial ribosomal proteins were no longer sufficient to secure the specific recognition of rRNA in a more complex archaeal cell.

Finally, apart from implications for evolutionary studies, finding NLS-type motifs in archaeal ribosomal proteins may contribute to biomedical research. This is because mutations in NLSs of eukaryotic ribosomal proteins are associated with Diamond-Blackfan anemia (DBA), a genetic blood disorder that is caused, among other factors, by defective ribosome biogenesis^42,43^. Particularly, mutation H11R in the NLS sequence of ribosomal protein uL3 was found in a DBA patient with defective ribosome biogenesis^44^. The fact the uL3 NLS-type motif is preserved in Archaea can make Archaea a unique and appealing model to study effects of NLSs mutations on ribosome biogenesis, because in Archaea the effect of these mutations on ribosome biogenesis can be studied independently of their effect on intracellular transport.

## Materials and Methods

### Comparison of protein sequences

The sequences of ribosomal proteins were retrieved from the Uniprot protein databank (https://www.uniprot.org) by using a script described in Ref.^45^. For species containing numerous copies of ribosomal proteins’ genes (for instance, *Saccharomyces cerevisiae* or *Arabidopsis thaliana*) only one isoform (isoform A or isoform 1) was used per species. For bacterial species in which several ribosomal proteins are encoded by two genes (corresponding to Zn-coordinating and Zn-free isoforms of ribosomal proteins), we only used the genes coding for Zn-coordinating isoforms. All the retrieved sequences are listed in (**Supplementary Data 1**). To create multiple sequence alignments we used the pre-compiled package of MAFFT (MAFFT-7.427) with default settings^46^.

To assess the conservation of NLS-type motifs in archaeal proteins, we calculated consensus sequences of NLS-type motifs for each of the four ribosomal proteins (uS12, uL3, uL15, and uL18) in each of the major archaeal branches (DAPNN, Euryarchaeota, TACK, and Asgard) by using multiple sequence alignments (MAFFT with default settings) of archaeal protein sequences listed in the (**Supplementary Data 1**). These consensus sequences were then compared with the corresponding consensus sequences of the eukaryotic NLSs. The NLS-consensus sequences were defined as follows: ^11-^RKLxxxRRxxRWxxxx-YKKRxxxxxxKxxP^-40^ for protein uS12 (residue numbers indicate their position in uS12 from *S. cerevisiae*, “x” designates any amino acid); ^3-^HRKxxxPRHxxxxxxPRKR^-21^ for protein uL3; ^4-^RxxKxRKxR^-12^ for protein uL15; and ^20-^FRRRRxxKxxY^-30^ for protein uL18. Sequence similarity shown in (**Fig. 2**) was calculated according to the AMAS definitions for the hierarchical analysis of residue conservation^47^ using Jalview^48^ (**Supplementary Data 2**).

### Comparison of ribosome structures

The ribosome structures were retrieved from the protein databank (https://www.rcsb.org) and were visualized and inspected by using PyMOL Molecular Graphics System (Version 2.0 Schrödinger, LLC.). Homologous ribosomal proteins were aligned by using “align” command to superpose Cα-atoms of conserved globular domains of homologous proteins or “pair_fit” command to superpose phosphate atoms of conserved rRNA segments in bacterial and archaeal ribosomes.

### Cloning of ribosomal proteins

The cDNA of human protein uS12 (NCBI Gene ID: 6228 and Uniport ID: P62266) and its N-terminally truncated mutant (deleted residues 1-41) were cloned into pEGFP-N1 vector (Promega) between KpnI and XhoI sites as described in Ref.19. To create chimeras between human uS12 and homologous proteins from *T. acidophilum* and *S. solfataricus*, the genomic sequence that corresponds to residues 1-41 of human uS12 was replaced by codon-optimized sequences from *T. acidophilum* (NCBI Gene ID: 1455748) and *S. solfataricus* (NCBI Gene ID: 38467843) corresponding to residues 1-37 of uS12 from *T. acidophilum* (Uniprot ID: Q9HLY2) or residues 1-42 from of uS12 from *S. solfataricus* (Uniport ID: P39573). The codon optimization for protein expression in human cells was done by using the codon optimization algorithm from the Integrated DNA Technologies (https://www.idtdna.com). The DNA fragments corresponding to the N-termini of archaeal uS12 were purchased from GenScript (https://www.genscript.com) and cloned into the *H. sapiens* uS12-pEGFP-N1 construct by using In-Fusion HD Cloning Plus (Takara).

### Cell lines, transfection and confocal microscopy

Nucleolar accumulation of eGFP-fused human ribosomal protein uS12 and its variants were examined in the human cell line HEK293T (ATCC, CCL11268). HEK293T cells were maintained in DMEM (Gibco) supplemented with 10% FBS (Gibco). Cells were plated on poly-L-lysine-coated glass cover slips at 70-80% confluence, and transfected with the respective plasmids encoding uS12 and its variants using Lipofectamine 2000 (Invitrogen), according to the manufacturers’ protocol.

Confocal fluorescent images were obtained by a Zeiss LSM 880 Airyscan NLO/FCS confocal microscope 48 h after transfection. Prior to imaging, cell samples were fixed with 4% paraformaldehyde, permeabilized with 0.1% triton X-100 and mounted using with ProLong^®^ Diamond Antifade Mountant (Invitrogen) with DAPI. For each eGFP-uS12 variant (and also for eGFP protein as a negative control), the snapshots were taken for six non-overlapping fields at 63x – for better data redundancy (**Supplementary Figure 1**). For each genetic construct, experiments were repeated three times.

## Authors Contributions

S.V.M. conceived the idea. S.V.M., C.C.T. and D.S. devised the study. S.V.M. and K.M. analyzed ribosome structures. H-S.K. and S.V.M. performed microscopy analysis. A.v.d.E. wrote scripts to analyze protein sequences. S.V.M. analyzed the sequences. All authors discussed the results and contributed to the final manuscript. S.V.M. and D.S. wrote the manuscript.

## Acknowdgments

We would like to thank members of Dieter Söll, Michael Rout and Marat Yusupov laboratories for valuable discussions at the early stage of this project. We also thank Richard Prum and Günter Wagner (Department of Ecology and Evolutionary Biology at Yale University) for critical feedback and stimulating discussions during preparation of the manuscript, and Vladimir Roudko (Icahn School of Medicine at Mount Sinai) for critical reading of the manuscript and Adam Bodley for editing. This work was supported by grants from the European Molecular Biology Organization (ASTF 434-2014 to S.M.) and from the National Institute of General Medical Sciences (R35GM122560 to D.S.).

