## Supplementary Figure 1 for "Archaeal ribosomal proteins possess nuclear localization signal-type motifs: implications for the origin of the cell nucleus"

**Figure S1 | Archaeal NLS-type motifs can functionally substitute NLS-signals of the eukaryotic ribosomal protein uS12 in human cells.** The panels show microscopic snapshots of eGFP fluorescence and fluorescence of the DNA-staining agent DAPI in the human cell line HEK293T.

A: eGFP – negative control (diffused localization between the nucleus and the cytoplasm)

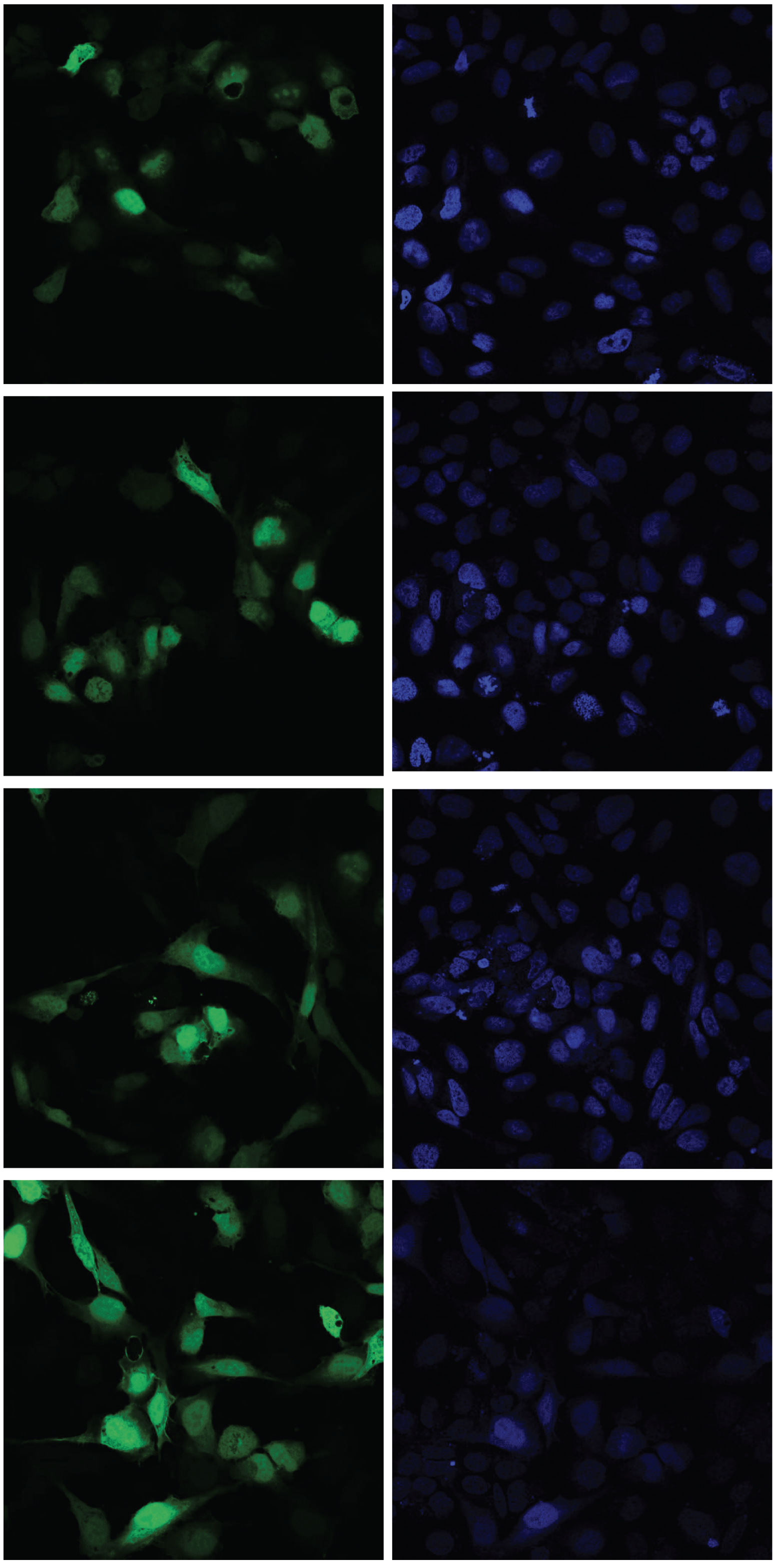

B: eGFP/human uS12 fusion (accumulated in the nucleoli)

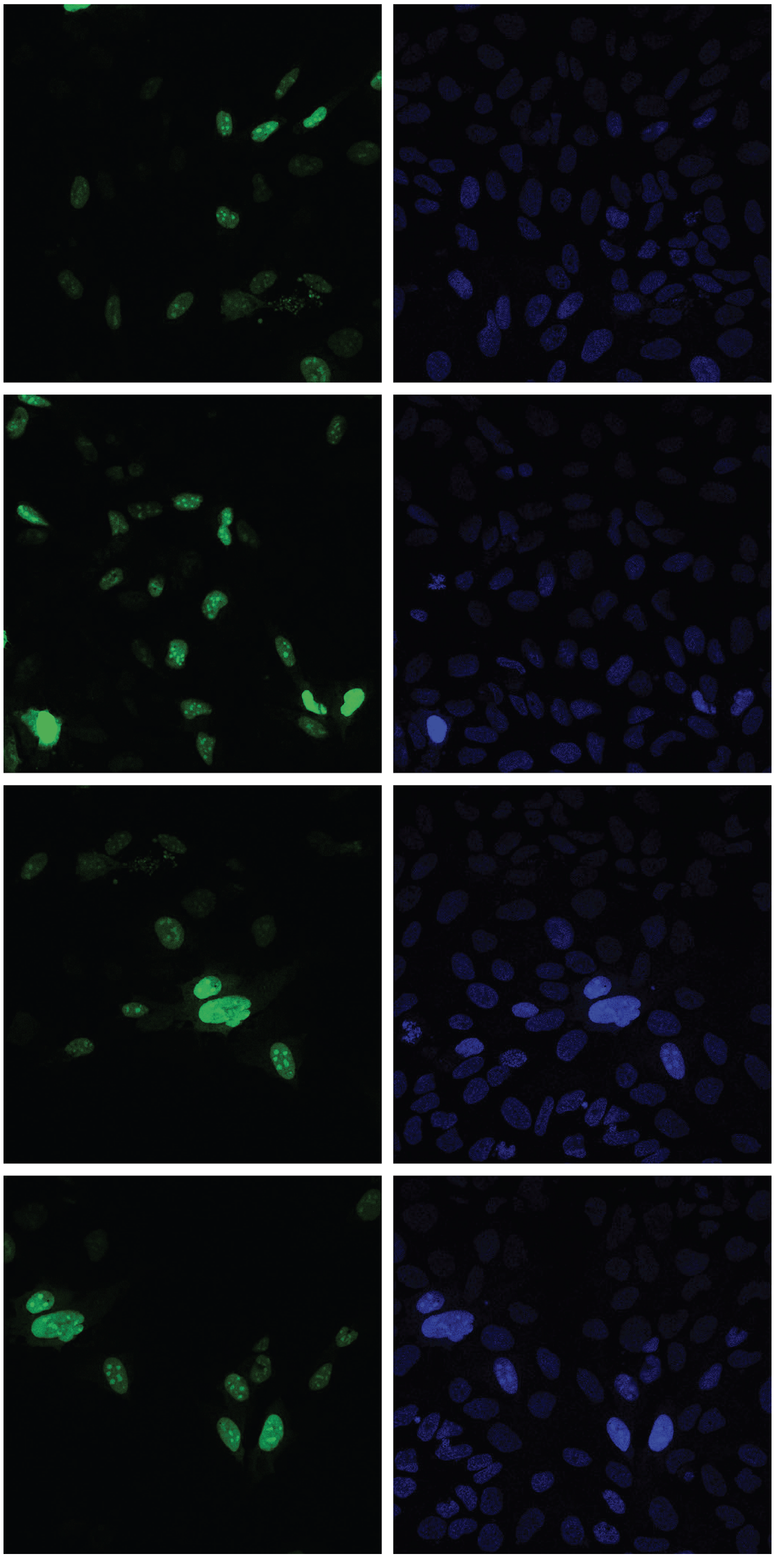

C: eGFP/human uS12 fusion with truncated NLS (excluded from the nucleoli)

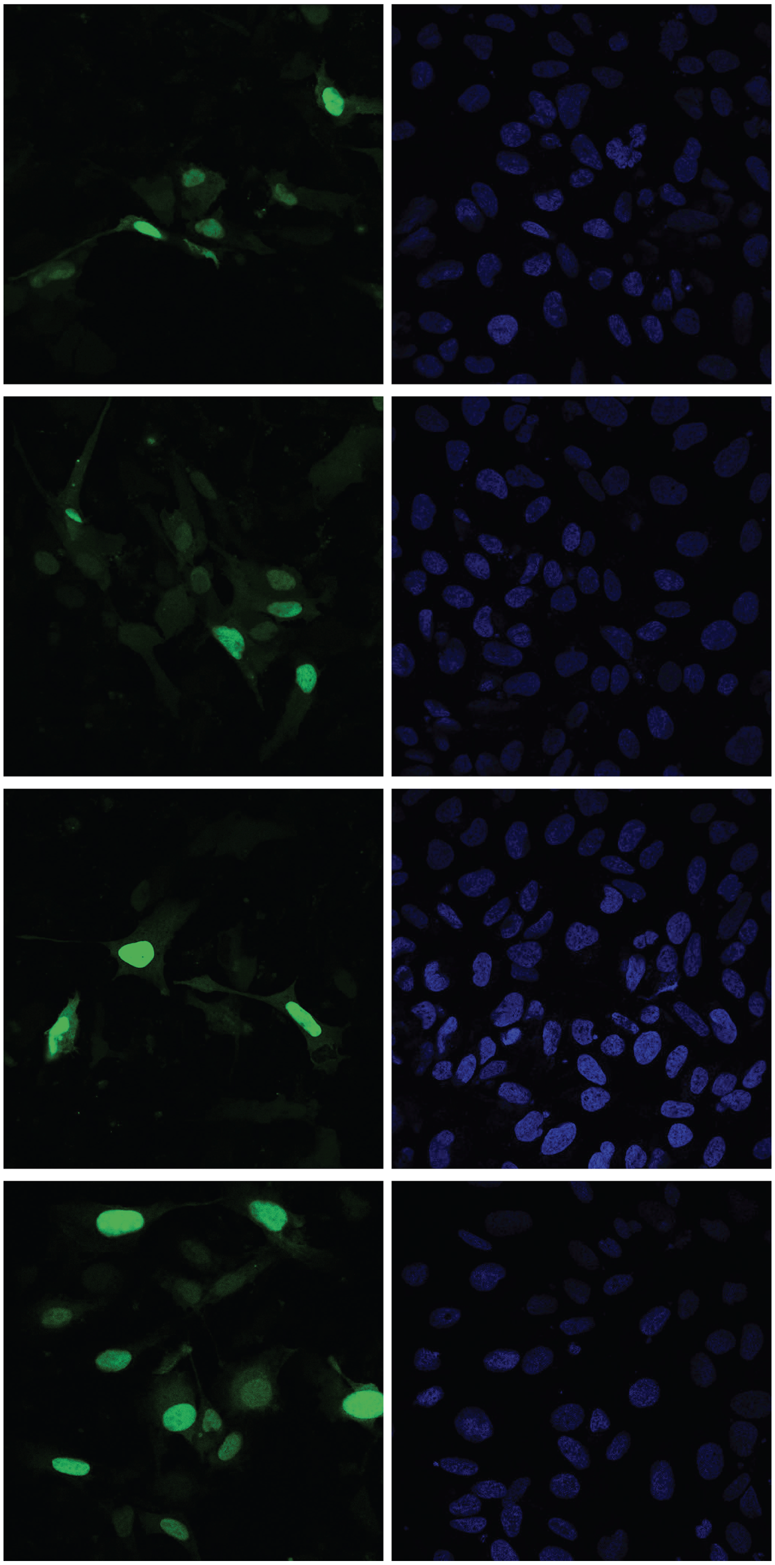

D: eGFP/human uS12 fusion with NLS-type segment from *S. solfataricus* uS12 (accumulated in the nucleoli)

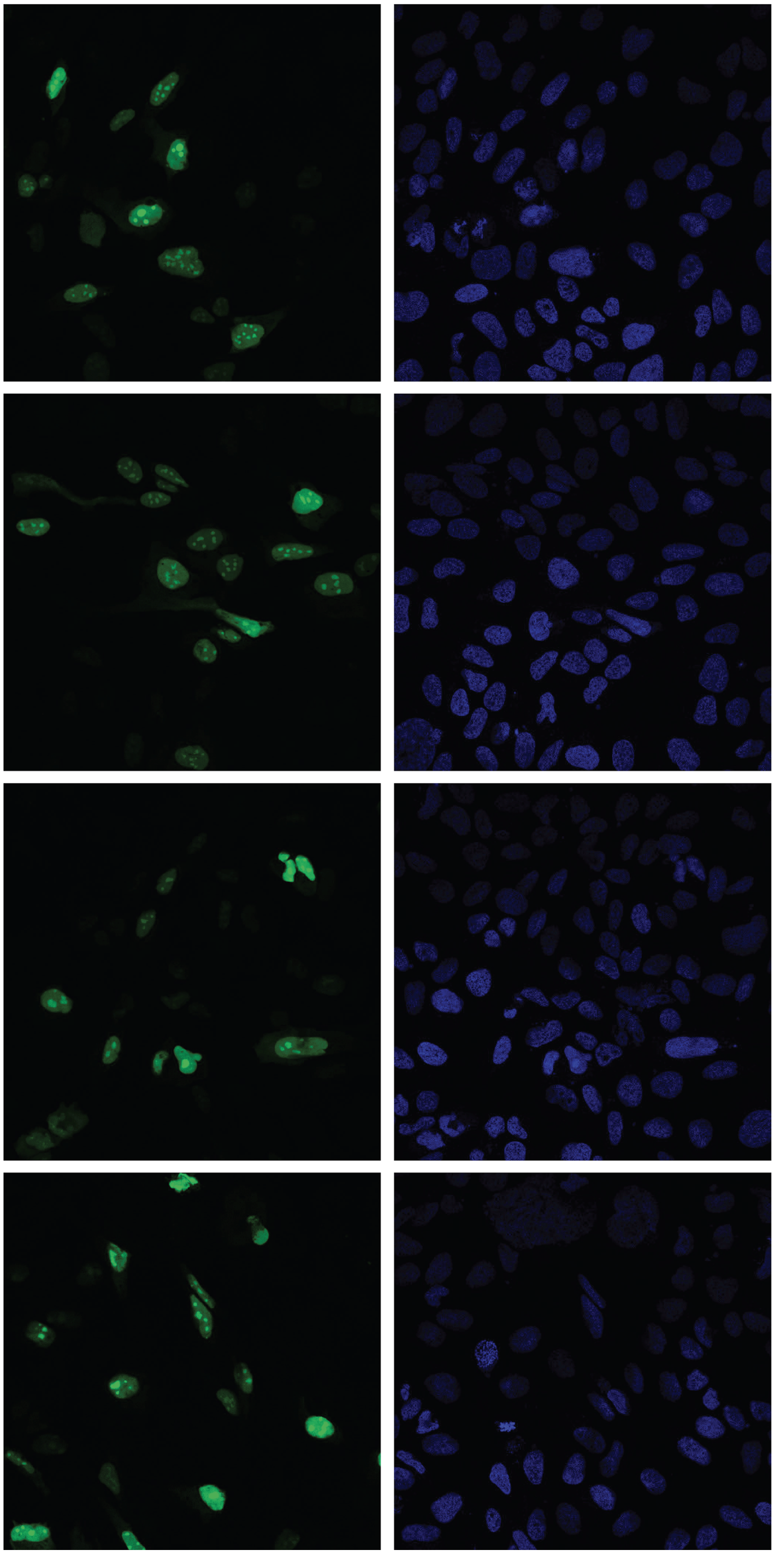

E: eGFP/human uS12 fusion with NLS-type segment from *T. acidophilum* uS12 (accumulated in the nucleoli)

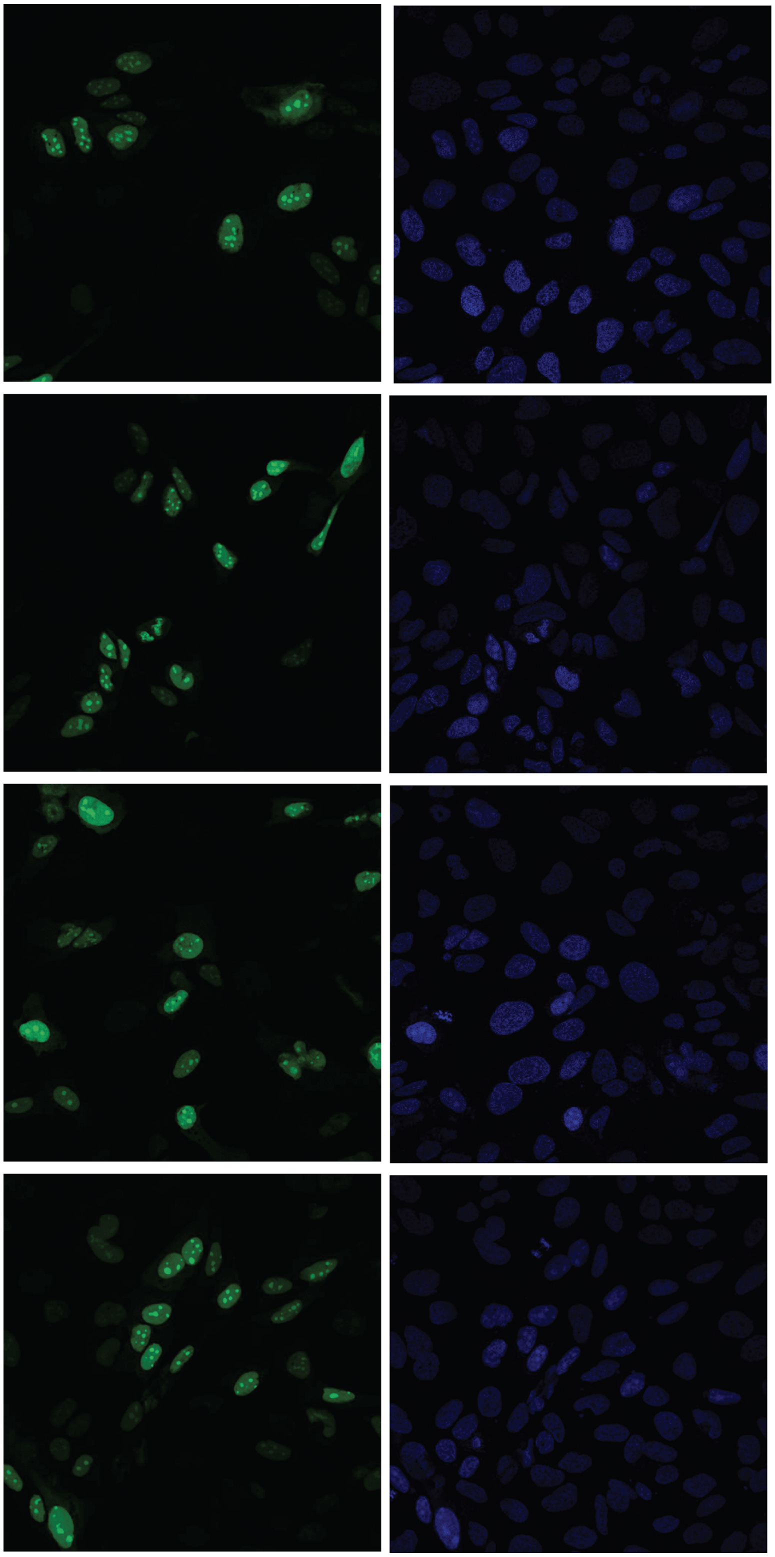
